# Accurate detection of non-proliferative diabetic retinopathy in optical coherence tomography images using convolutional neural networks

**DOI:** 10.1101/667865

**Authors:** Mohammed Ghazal, Samr Ali, Ali Mahmoud, Ahmed Shalaby, Ayman El-Baz

**Affiliations:** Department of Electrical and Computer Engineering, Abu Dhabi University, Abu Dhabi, United Arab Emirates; Department of Electrical and Computer Engineering, Concordia University, Montreal, Canada; BioImaging Laboratory, Department of Bioengineering, University of Louisville, Kentucky, United States of America

**Author notes:** Corresponding author (MG).

## Abstract

Diabetic retinopathy (DR) is a disease that forms as a complication of diabetes, It is particularly dangerous since it often goes unnoticed and can lead to blindness if not detected early. Despite the clear importance and urgency of such an illness, there is no precise system for the early detection of DR so far. Fortunately, such system could be achieved using deep learning including convolutional neural networks (CNNs), which gained momentum in the field of medical imaging due to its capability of being effectively integrated into various systems in a manner that significantly improves the performance. This paper proposes a computer aided diagnostic (CAD) system for the early detection of non-proliferative DR (NPDR) using CNNs. The proposed system is developed for the optical coherence tomography (OCT) imaging modality. Throughout this paper, all aspects of deployment of the proposed system are studied starting from the preprocessing stage required to extract input data to train the CNN without resizing the image, to the use of transfer learning principals and how best to combine features in order to optimize performance. A novel patch extraction framework for preprocessing is presented, followed by fovea detection algorithm, in addition to investigating the various CNN parameters for optimal deployment. Optimum CNN parameters and promising results are achieved. To the best of our knowledge, this is the first CNN-based DR early detection CAD system for OCT images. It achieves a promising accuracy of 94% with transfer learning.

## Introduction

Nowadays, ophthalmologists are capable of determining diseases with the utilization of computer aided diagnostic (CAD) systems for accurate diagnosis, in contrast to the traditional form of visual interpretation and observation. Although CAD systems have recently emerged in the medical field, there is a continuous flux of interest in the development of such systems due to their capability of improving the medical services provided to the community in terms of accuracy and reliability in the diagnosis of diseases. Meanwhile, machine learning is paving the way for breakthroughs in the different areas of medical imaging such as in classification [1], segmentation [2], disease detection [3], and image registration [4]. The application of deep learning, a subset of machine learning algorithms, has made tremendous impact in the area of medical image processing research [5, 6]. Deep learning is the latest emerging machine learning lead technology in computer vision and image processing domains, particularly convolutional neural networks (CNNs) [7]. They are especially powerful in solving problems that are computationally difficult or with a high error rate such as medical image recognition with outstanding performance results [8]. For this reason, we got inspired to use CNNs for the early detection of one of the most serious ophthalmological problems, which is the diabetic retinopathy (DR).

Blindness resulting from diabetes is turning to be an increasingly alarming issue, which is due to the associated eye disease: DR. Such disease which develops as a complication of diabetes, particularly type II [9, 10] occurs specifically from the chronic high levels of sugar in the blood associated with swelling and damage of the tiny retinal blood vessels in the eye [11–13]. This leads to distortion of the vision followed by scarring of the retina in advanced stages, and finally consequent blindness [12]. It is worth mentioning that DR is one of the leading causes of blindness in adults [14, 15]. This problem is further amplified by the fact that 75% of the diabetic patients are not aware of the eye complications that they may be experiencing [9]. To prevent such a ramification, it is paramount to diagnose DR as soon as possible. Early detection and intervention can slow down the process and halt it completely [16, 17], which in turn protects the vision of the patient [18]. However, despite the significance of this matter and the notable prevalence rise of diabetes, a precise procedure to detect early retinal changes for DR prevention is absent [13, 17].

DR has mainly two distinct classes: proliferative DR (PDR) and non-proliferative DR (NPDR) [6]. NPDR is characterized by the presence of damaged blood vessels in the retina in addition to fluid leakage, which results in the retina swelling and wetness. For the case of PDR, multiple regions of the retina are affected by the appearance of new abnormal blood vessels, which makes it a severe and advanced DR stage. The work presented in this paper is limited only to NPDR.

One of the ophthalmology CAD imaging modalities used for early DR detection is fundus images [19, 20]. This is a particularly active area for research in applications of automatic detection because while fundus imaging are convenient in terms of its imaging principals that correspond to ophthalmoscopy, their interpretations requires a highly trained ophthalmologist; hence it is expensive [21]. Related work on fundus imaging for early detection of DR includes automatic micro-aneurysms detection trainable system [22], rendering a 65% sensitivity at 27 false positives per image by supposition testing. Another system, though dependent on the existence of the indicators, is able to detect hard and soft exudates [23]. On the other hand, in [24], Pachiyappan et al. use a combination of filtering, morphological processing, and thresholding for DR macular abnormalities detection, while in [25], automatic extraction of retinal vasculature was performed in order to obtain the blood vessels network. Similar feature-based for fundus imaging have also been reviewed in [26]. Recently, CNNs have also been used for exudates detection of diabetic subjects in fundus images [27].

Nonetheless, another medical imaging modality that may be employed in the early detection of DR is optical coherence tomography (OCT) [28], which is useful because it facilitates retinal morphology evaluation to microscopic resolution [29]. In turn, various retinal abnormalities including glaucoma, macular degeneration, and diabetic macular edema may be diagnosed in a non-invasive manner. Compared to fundus imaging, OCT is more favorable because it supports quantitative evaluations as it capable of capturing depth, in addition to its lower cost, and its ability to allow human bias free monitoring of changes [30]. However, OCT is relatively unexplored in comparison to fundus images in terms of early detection of DR. As such, the proposed system investigates early detection of DR in OCT images, which is principally performed with the aid of CNNs.

Hence, this paper presents the optimal CNN architecture for early detection of DR through the exploration of various CNN configurations and parameters. In particular, the following items are studied: 1) the effect of transfer learning on improving the performance of the proposed CAD system, given the scarcity of the data; 2) the effect of fusing CNNs retrained with different datasets on the overall system performance; 3) the depth of CNN layers required to extract features to train the final classifier used for data fusion; and 4) the OCT layer required to be segmented in order to be used for the y-coordinate axis alignment of the extracted patches for optimum results. The rest of this paper is organized in three sections as follows: a section that presents the materials and methods, followed by a section that discusses the experimental results, and finally the conclusion is given in the last section.

## Materials and methods

A simplified block diagram of the proposed CAD system for early detection of DR in OCT images using CNNs is shown in Fig 1, and an overview of the system is shown in Fig 2. In contrast to conventional feature extraction methods, using CNNs effectively classifies normal and DR images without the need for features that are designed manually. The proposed system is composed of a preprocessing stage where appropriate sized patches are detected and extracted, a CNN training stage, and finally investigation of various fusion or combination schema for improved performance of the proposed system. The preprocessing stage starts with segmentation of the original OCT scan into twelve different layers with the application of an unsupervised parametric mixture model and Markov Gibbs Random Fields [31]. The location of the fovea is also simultaneously detected. The results of the prior two stages are then fed for the positioning and extraction of the appropriate patches as per the schema of the proposed system.

**Fig 1.**
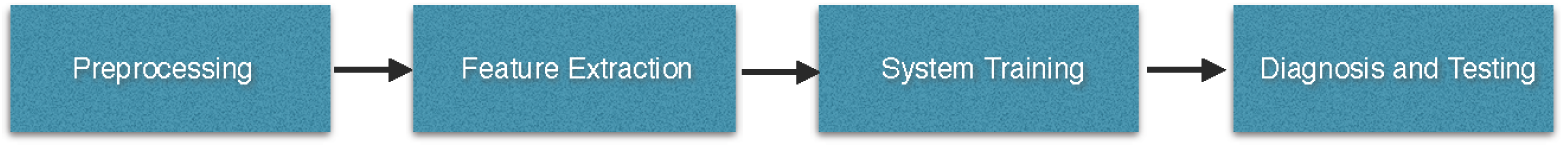
Simplified block diagram of the proposed CAD system for early detection of DR in OCT images using CNNs.

**Fig 2.**
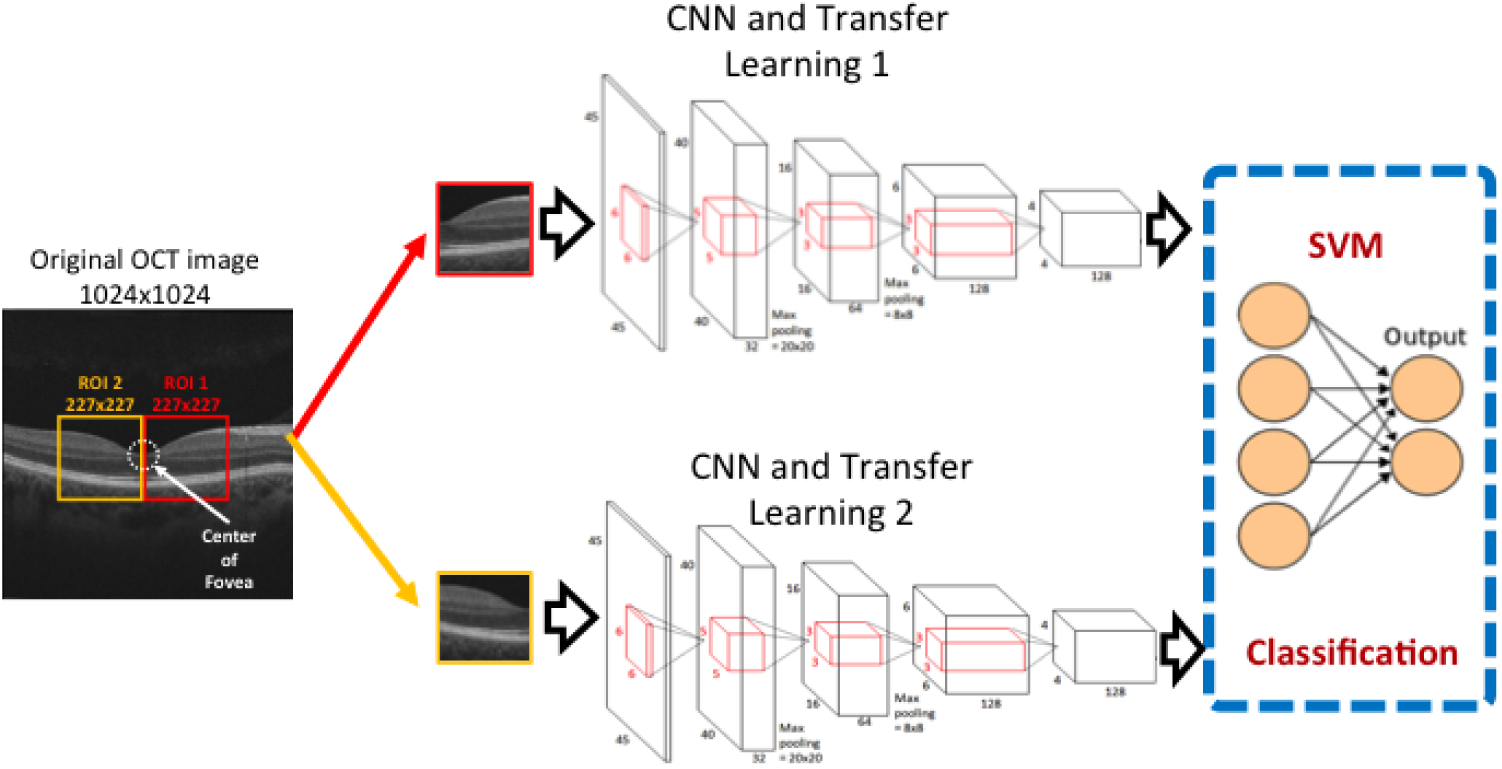
Overview of the proposed CAD system for early detection of DR in OCT images using CNNs.

The extracted patches are then fed into the CNNs of the proposed system according to the configuration at which the system has been set. The output from the CNNs is then analyzed if it is determined to be the final output of the proposed CAD system; otherwise, it is fed into a classifier for appropriate training for combination of the data to obtain the final result. We evaluate the overall performance of the system with different metrics and the utilization of 5-fold cross validation technique. This allows us to infer meaningful conclusions and develop the optimum CAD system for early detection of DR with the use of CNNs.

### OCT dataset

Patients with and without diabetes mellitus were enrolled at the Kentucky Lions Eye Center at University of Louisville from between June 2015 and December 2015 (total of 52 subjects with 26 patients for each class with IRB Number: 18.0010). Informed consent (or assent) was obtained for each participant. Exclusion critiera included history of retinal pathology, including diabetes-related, and severe myopia, defined as refractive error ≥ −6.0 diopters. In all, *n* subjects were enrolled, *n* of whom had diabetes, ranging in age from *n* to *n* years.

Data used for training and testing of the CAD system were obtained using a clinical OCT scanner, Cirrus HD-OCT 5000 (Carl Zeiss Meditec, Dublin, California). B-scans were obtained over a 21-line raster across the macula of both eyes. For each eye, a single b-scan, passing through the fovea, was selected for analysis. Images were 1024 ×1024 pixels, 8 bit grayscale, capturing an optical slice 2 mm deep and 9 mm from side to side (nasal-temporal).

### Preprocessing and retinal layer segmentation

The proposed DR early detection system first constitutes of a preprocessing stage, as illustrated in Fig 3, where we extract the inputs to be fed to the CNN as appropriate. The main reason behind such a stage is the disagreement between the dimensional size of the original OCT scans that are the input image data and that of the input layer for the pretrained CNN in the methodology; i.e. the AlexNet CNN as shown in Fig 4. The pre-trained version of the CNN for the application of transfer learning is trained on a subset of the large-scale ImageNet image database [32]. That is 1000 object categories and 1.2 million real-life images for training. The preprocessing stage also eliminates unimportant information; hence, improving the speed and efficiency of the system.

**Fig 3.**
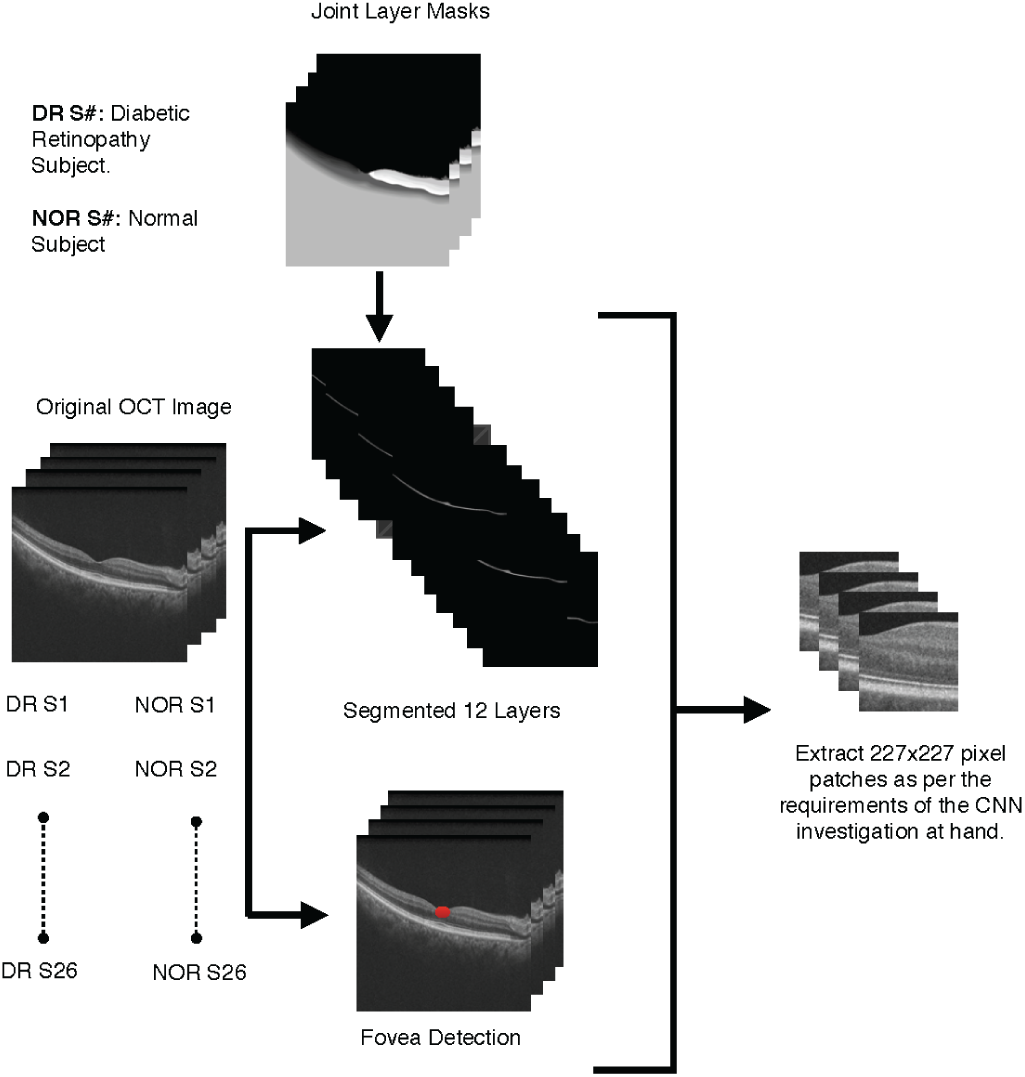
Preprocessing stage of the proposed CAD system.

**Fig 4.**
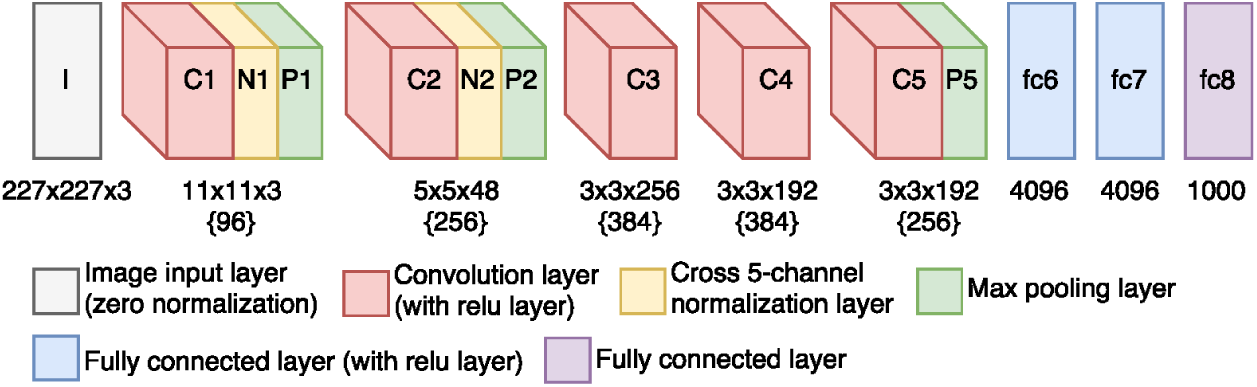
Pre-trained AlexNet CNN model architecture.

Preprocessing for the proposed algorithm involves rough segmentation of the retina proper from the rest of the image, and identification of the fovea. The appearance of the retina in an OCT b-scan reveals approximately 12 bands of greater or lesser reflectivity as shown in Fig. 5. Histological studies have correlated these bands with the layers of the retina, proceeding from the vitreous body to the choroid: 1. nerve fiber layer (NFL), 2. ganglion cell layer (GCL), 3. inner plexiform layer (IPL), 4. inner nuclear layer (INL), 5. outer plexiform layer (OPL), 6. outer nuclear layer (ONL),7. external limiting membrane (ELM), 8. myoid zone (MZ), 9. ellipsoid zone (EZ),10. outer segments of the photoreceptors (OPR), 11. interdigitation zone (IZ), and 12.the retinal pigment epithelium (RPE).

**Fig 5.**
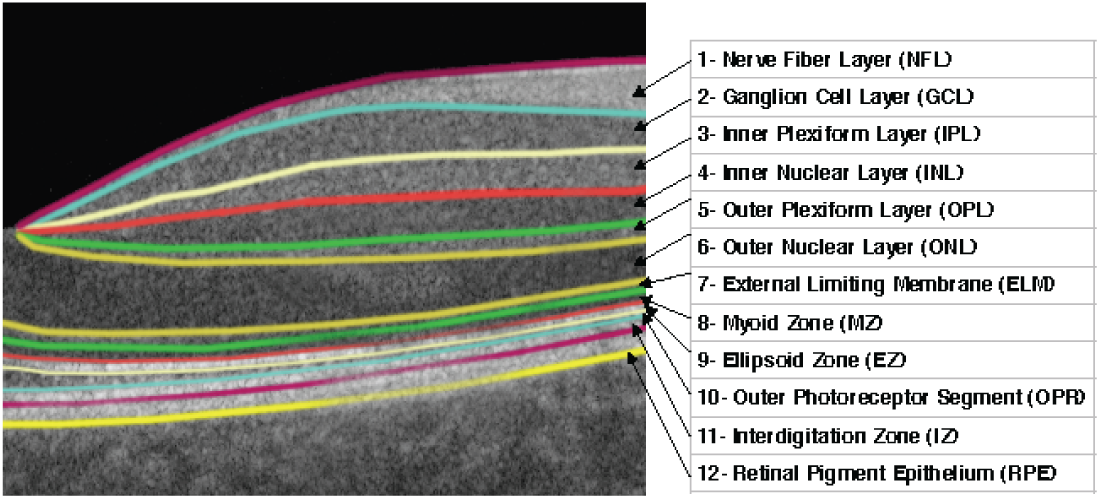
OCT retinal layers segmentation.

As shown in Fig 5, at the fovea the vitreous body is nearly adjacent to the ONL (layer 6), while layers 1–5 all but vanish; surrounding the fovea is the foveal rim, where layers 1–5 are thickest. This is the structure used to guide the patch extraction procedure for consistent representation of the retina across different OCT images. Patches were selected from either side of the fovea and oriented to align with the retinal layers. Consequently, the amount of extracted background (vitreous or choroid) is minimized, and feature extraction should be independent of any peculiarities (e.g. slight tilt or off-center) of a given OCT scan.

The fovea coordinate localization starts off with applying a median filter in order to remove any impulsive noise. This is then followed by the “à trous” algorithm [33] that decomposes each scan into scale-space components of coarser and finer detail by undecimated wavelet transform as per the aforementioned details. Edge detection in scale space allows for easy identification of high contrast boundaries in the OCT image: vitreous-NFL, MZ-EZ, and RPE-choroid. Contours are first detected as local gradient maxima in the appropriate wavelet component, then smoothed using adaptive spline smoothing. Considerations of typical retina structure, above, lead to identification of the fovea with the point on the vitreous-NFL boundary at minimum distance from the MZ-EZ boundary. When computing these distances, it is important to correct for the non-square pixel aspect ratio of typical OCT scanners. The preprocessing algorithm is described in detail in [31]. Finally, it is noteworthy to mention that each of the grayscale images was concatenated as three different channels for the required 3-channel input of the AlexNet.

### Feature extraction and diagnosis

Upon the detection of the fovea, the required patches’ locations along the x-coordinate axis is computed appropriately, i.e. with the origin at the fovea. These calculated points are used for extracting the corresponding vertical slice from the segmentation mask of the layer that we would be centering the patch extraction at for the y-coordinate axis. The sum the values of the pixels of these extracted slices along the x-axis is then considered to take into account any orientation or skewness display of the OCT scan. It acts as an efficient measure where if it is greater than zero then the corresponding row contains a significant part of the layer. The resultant of the final algorithm of the preprocessing stage as shown in Fig 6 is a representative patch, i.e. extracted image from the original OCT, that is in a matching size to the input layer of the CNN.

**Fig 6.**
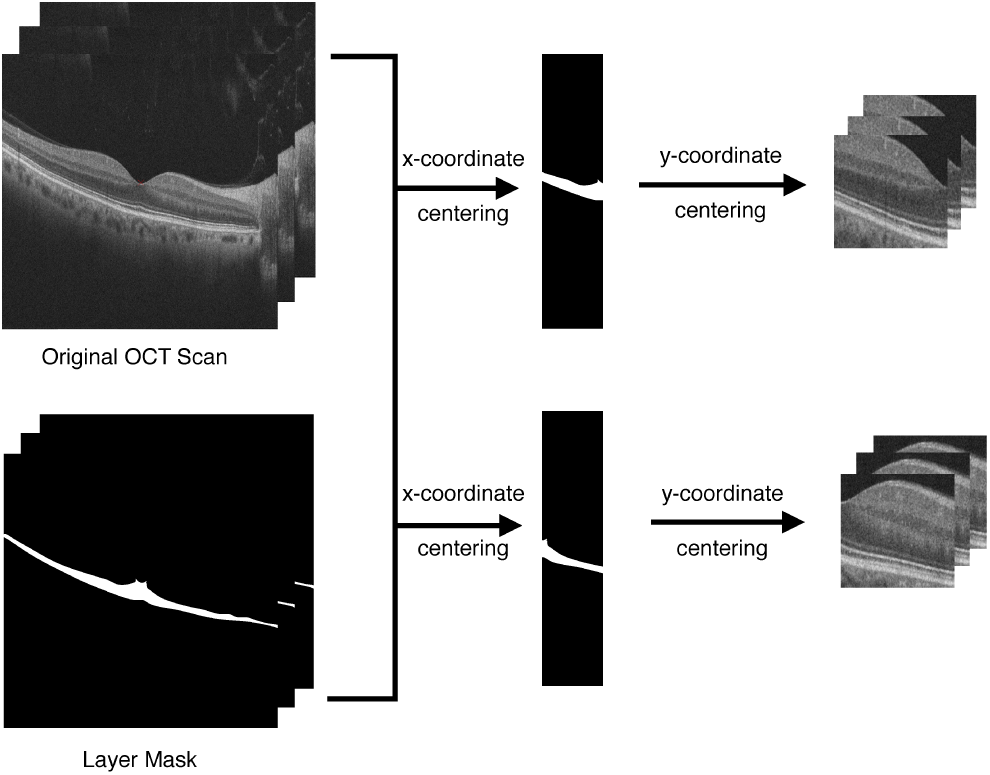
Patch extraction framework.

In order to find out the optimum parameters for the proposed system, various investigations were carried out, the first of which is whether the application of transfer learning improves the accuracy of the algorithm rather than just training the network from scratch. This is also made in accordance to the fact that the dataset size is relatively small. As such, training of two AlexNet CNNs is performed; one which is pre-trained and another which is not, with the normal and DR training dataset upon its preprocessing.

Patches extracted to the temporal side of the fovea were fed into their own AlexNet CNN, one of which was pretrained on ImageNet and another which had not been. The pretraining is used only as far as layer *pool5* (Fig. 4) for the activations of the hidden layers, while subsequent layers of the AlexNet were treated identically for the comparison between the use and non-use of transfer learning. Patches extracted to the nasal side of the fovea were likewise input to two AlexNet CNN, one of which was pre-trained, for carrying out the same comparison.

It was also tested whether improved accuracy would result if fusion of various input data. This experiment was carried out through different means for extracting input data and building a corresponding appropriate CNN configuration for testing and validation of the investigation. The first use case is the configuration of the best results concluded from the first experimentation. That is hypothetically the two pretrained AlexNet CNNs that are fed the input images as the patches extracted to the left and to the right of the fovea; i.e. the experiment is hypothesized to show that transfer learning does improve the results particularly with small datasets. Fovea detection preprocessing is then performed for each of the OCT scans or images in order to extract a patch where the fovea coordinates are the center of the patch. That is the extraction process of the patch assumes the fovea coordinates to be its center across both x-coordinates and y-coordinates instead of just the x-coordinates and relying on the segmented layer (6 or ONL) chosen as per the other extraction scheme. Hence, successfully combining the information from the left and right patch to be fed to a single AlexNet CNN and test the accuracy.

Furthermore, information combination of the patches extracted to the left and right of the fovea in another way which aims to further improve the results by increasing the amount of information fed to the network or the AlexNet CNN in investigated. That is whether the combination of the results of the CNNs that are trained with the patches extracted to the left and right of the fovea can improve the overall algorithm’s performance. This is carried out by the extraction of the features at the bottleneck stage and then the training of a two-class support vector machine (SVM) classifier, a machine learning algorithm used in classification problems [34], with a fast stochastic gradient descent solver for the final classification results.

Hence, the proposed overall CNN configuration starts with the fovea detection in order to locate it as well as the layer segmentation to acquire the resultant segmented layer 6, i.e. ONL layer, of each of the scans that we are keeping as a constant parameter for the experiments carried out. However, it is noteworthy to mention that a different experimentation is carried out where all other parameters are set but the layer at which we center the patch extraction process at is varied in terms of the y-coordinates to find the layer that achieves the optimum results and hence further improves the algorithm. This entire CNN setup is shown in Fig 7.

**Fig 7.**
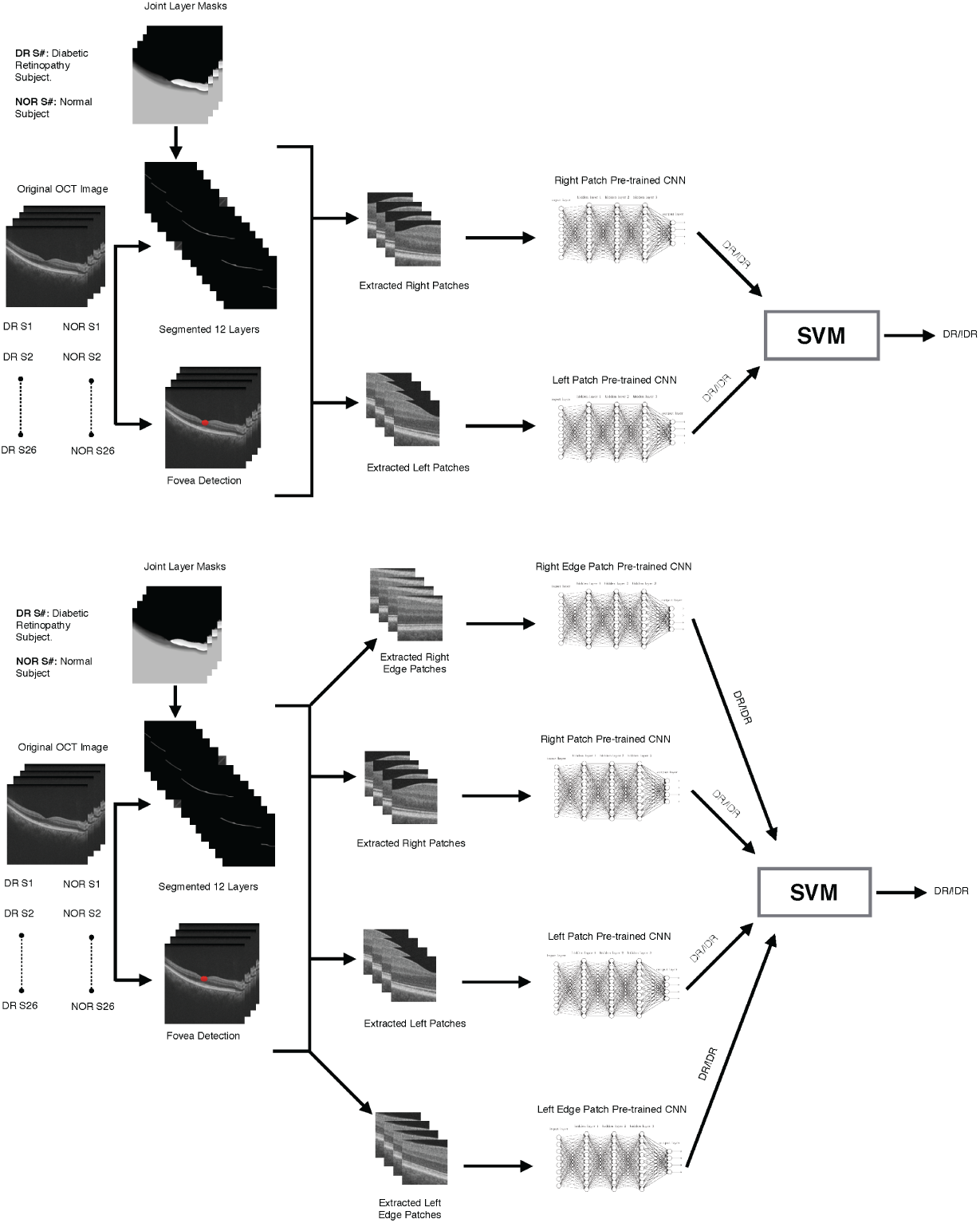
Investigation of deployment of various data fusion schemes in the proposed system with transfer learning. A: Patches extracted to the left and right of the retina combined convolutional neural networks together. B: Patches extracted to the left and right of the retina, and left and right edge patches combined convolutional neural networks.

Another investigation is carried out in order to find out the optimum parameters for the best performance of the algorithm is the layer at which we carry out transfer learning at by change of extracting the features at the following layers for comparison of the performance results: (i) one layer above the bottleneck features which is relu6 layer, *f{c6}*; (ii) the layer with the bottleneck features which is pool5 layer, *{P5}*; (iii) one layer before the bottleneck features which is relu5 layer, *{C5}*; and (iv) two layers before the bottleneck features which is relu4 layer, *{C4}* In this investigation, both the results in terms of accuracy and the computation expense for an overall performance evaluation are taken into consideration.

The final investigation finds out the best OCT segmented layer to center the patch extracted process at building on the results from the above investigations. The layers that are experimented on are all around the location of the fovea which are set to be: (i) layer 5 or the OPL; (ii) layer 6 or the ONL; (iii) layer 7 or the ELM; (iv) layer 8 or the MZ; and (v) layer 9 or the EZ. Samples of such extracted patches are shown in Fig 8 for DR OCTs.

**Fig 8.**
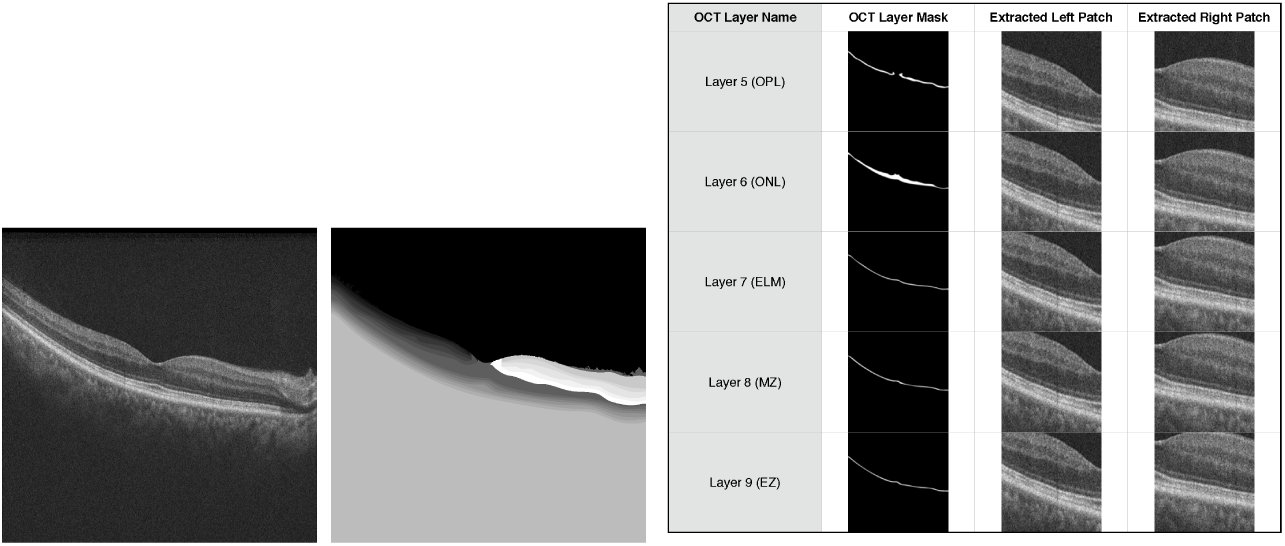
DR sample patches centered with different OCT layers. A: Original OCT scan from a subject with DR. B: OCT scan corresponding joint layers mask for segmentation. C: Patches extracted to the left and right of the retina from a subject with DR centered across the y-coordinate axis with different OCT layers.

### Validation

Fivefold cross validation was used for the evaluation of each of the investigations of the CNN CAD system for early detection of DR. This is a particular case of *k*-fold cross validation, where *k* training runs are performed, each time leaving out a fraction 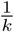 of the data for subsequent testing. In this way every available observation (OCT scan) is used for both training and (exactly once) for testing. This is in contrast to the traditional hold-out method where a fixed proportion of the data are set aside from the beginning of the experiment to be used only for validation. Cross validation has the advantage of making the most of a limited amount of data, but is known to produce biased estimates of system performance.

The performance metrics used were accuracy (*α*), error rate (*β*), specificity (*χ*), precision (PPV), and recall (TPR). If TP is the number of correctly classified diabetes cases in a particular run of cross validation, TN is the number of correctly classified non-diabetic cases, and P = TP + FP and N = TN + FN are the total number of diabetes and non-diabetic cases in the test data, then these metrics are defined as

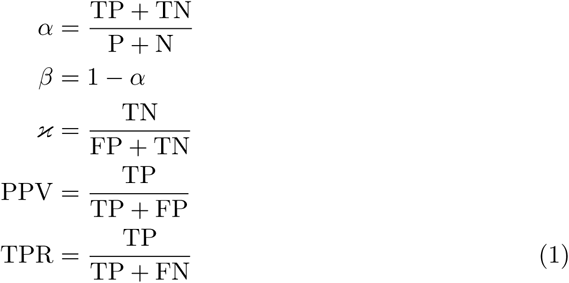

### Experimental results and discussion

Shown in Table 1 are the comparative results of the proposed investigations. In addition to the aforementioned performance metrics, the respective standard deviations across the folds are noted. The standard deviation of the error metric corresponds to that of the accuracy as they are complementary measures. First, when investigating the effect of transfer learning application on the proposed system for a CNN which is fed only patches extracted to the left and to the right of the fovea as input, a clear increase in performance across all of the measures is observed. Second, comparison of the performance results of training the proposed system with independent patches versus various combination schemas such as patches extracted with the fovea at their center, or using a classifier to combine two or four extracted patches shows that combining two or four patches result in the highest performance metrics across the board. However, it can also be observed that the time taken for the four patches approach is almost four times the one for the two patches, with no improvement in any of the metrics. As such, only the two patches approach is accounted for in the next investigation of finding the optimum CNN layer to extract the features from. It can be derived from the summarization in the table that the bottleneck features represent the best choice in terms of balance between the run time required, i.e. computation complexity and the accuracy. Finally, to choose the OCT layer to align the extraction of the input patches at, we find out that layer 6 is the best given that it reaches the highest performance metrics. That is though other layers achieve similar levels of accuracy, error rate, specificity, precision, and recall, they all require more time to train. As such, the final design choices for the proposed system is the CNN retrained with two patches extracted to the right and the left of the fovea centered with OCT layer 6 across the y-coordinate axis. The features are extracted from *pool5* CNN layer and transfer learning is applied in the deployment of the proposed system.

**Table 1.** Summary of proposed system findings.

|  | Accuracy | Accuracy Std | Error Rate | Specificity | Specificity Std | Precision | Precision Std | Recall | Recall Std | Average Testing Time (s) |
| --- | --- | --- | --- | --- | --- | --- | --- | --- | --- | --- |
| Right Alone (No transfer learning) (pool5 - layer6) - 128 batches | 52.00% | 0.08940 | 48.00% | 41.43% | 0.09620 | 64.44% | 0.20560 | 83.81% | 0.23480 | 0.403820 |
| Right Alone (No transfer learning) (pool5 - layer6) - 64 batches | 56.00% | 0.08940 | 44.00% | 62.50% | 0.21650 | 65.11% | 0.21150 | 88.67% | 0.17580 | 0.492000 |
| Right Alone (No transfer learning) (pool5 - layer6) - 32 batches | 54.00% | 0.05480 | 46.00% | 42.86% | 0.04120 | 66.67% | 0.20410 | 84.29% | 0.20450 | 0.486904 |
| Left Alone (No transfer learning) (pool5 - layer6) - 128 batches | 44.00% | 0.05480 | 56.00% | 36.67% | 0.21010 | 57.83% | 0.22600 | 71.79% | 0.20220 | 0.386900 |
| Left Alone (No transfer learning) (pool5 - layer6) - 64 batches | 56.00% | 0.15170 | 44.00% | 59.68% | 0.22980 | 62.00% | 0.21680 | 89.33% | 0.15350 | 0.492300 |
| Left Alone (No transfer learning) (pool5 - layer6) - 32 batches | 50.00% | 0.00000 | 50.00% | 30.00% | 0.00000 | 76.79% | 0.22520 | 66.79% | 0.20660 | 0.4794120 |
| Right Alone (Transfer learning) (pool5 - layer6) - 128 batches | 80.00% | 0.07070 | 20.00% | 77.78% | 0.10190 | 80.00% | 0.07070 | 100.00% | 0.00000 | 0.394520 |
| Right Alone (Transfer learning) (pool5 - layer6) - 64 batches | 86.00% | 0.11400 | 14.00% | 80.12% | 0.14160 | 86.00% | 0.11400 | 100.00% | 0.00000 | 0.160400 |
| Right Alone (Transfer learning) (pool5 - layer6) - 32 batches | 92.00% | 0.08370 | 8.00% | 89.33% | 0.09830 | 93.78% | 0.05700 | 97.78% | 0.05640 | 0.156400 |
| Left Alone (Transfer learning) (pool5 - layer6) - 128 batches | 74.00% | 0.04470 | 26.00% | 73.00% | 0.06190 | 80.78% | 0.06690 | 90.78% | 0.04970 | 0.378820 |
| Left Alone (Transfer learning) (pool5 - layer6) - 64 batches | 76.00% | 0.08940 | 24.00% | 68.45% | 0.09170 | 76.00% | 0.08940 | 100.00% | 0.00000 | 0.15900 |
| Left Alone (Transfer learning) (pool5 - layer6) - 32 batches | 66.00% | 0.08940 | 34.00% | 66.72% | 0.18840 | 73.00% | 0.16430 | 91.00% | 0.12450 | 0.15690 |
| Center (No transfer learning) (pool5 - layer6) - 128 batches | 52.00% | 0.10950 | 48.00% | 58.89% | 0.23110 | 58.89% | 0.23110 | 90.00% | 0.14140 | 0.487400 |
| Center (No transfer learning) (pool5 - layer6) - 64 batches | 52.00% | 0.04470 | 48.00% | 23.33% | 0.11790 | 87.14% | 0.21670 | 63.33% | 0.21730 | 0.498400 |
| Center (No transfer learning) (pool5 - layer6) - 32 batches | 56.00% | 0.08940 | 44.00% | 51.11% | 0.24220 | 76.29% | 0.22940 | 76.50% | 0.22610 | 0.445120 |
| Center (Transfer learning) (pool5 - layer6) - 128 batches | 74.00% | 0.15170 | 26.00% | 69.45% | 0.13550 | 82.44% | 0.08070 | 88.89% | 0.19250 | 0.467270 |
| Center (Transfer learning) (pool5 - layer6) - 64 batches | 78.00% | 0.08370 | 22.00% | 72.79% | 0.08780 | 81.33% | 0.08290 | 95.28% | 0.06480 | 0.456780 |
| Center (Transfer learning) (pool5 - layer6) - 32 batches | 80.00% | 0.10950 | 20.00% | 80.29% | 0.13470 | 87.33% | 0.08330 | 91.28% | 0.15350 | 0.431520 |
| Resized Image (Transfer learning) (pool5) - 32 batches | 78.00% | 0.13040 | 22.00% | 71.23% | 0.12400 | 78.00% | 0.13040 | 100.00% | 0.00000 | 0.406880 |
| Four Patches (Transfer learning) (pool5 - layer6) - 32 batches | 94.00% | 0.05480 | 6.00% | 88.00% | 0.10950 | 90.00% | 0.09130 | 100.00% | 0.00000 | 0.420160 |
| Two Patches (Transfer learning) (pool5 - layer6) - 32 batches | 94.00% | 0.05480 | 6.00% | 88.00% | 0.10950 | 90.00% | 0.09130 | 100.00% | 0.00000 | 0.420160 |
| Two Patches (Transfer learning) (relu6 - layer6) - 32 batches | 84.00% | 0.05480 | 16.00% | 72.00% | 0.17890 | 79.52% | 0.12250 | 96.00% | 0.08940 | 0.429200 |
| Two Patches (Transfer learning) (relu5 - layer6) - 32 batches | 94.00% | 0.05480 | 6.00% | 88.00% | 0.10950 | 90.00% | 0.09130 | 100.00% | 0.00000 | 0.420160 |
| Two Patches (Transfer learning) (relu4 - layer6) - 32 batches | 92.00% | 0.04470 | 8.00% | 84.00% | 0.08940 | 86.67% | 0.07450 | 100.00% | 0.00000 | 0.406120 |
| Two Patches (Transfer learning) (pool5 - layer5) - 32 batches | 88.00% | 0.04470 | 12.00% | 88.00% | 0.10950 | 90.00% | 0.09130 | 88.00% | 0.17890 | 0.431940 |
| Two Patches (Transfer learning) (pool5 - layer7) - 32 batches | 94.00% | 0.05480 | 6.00% | 88.00% | 0.10950 | 90.00% | 0.09130 | 100.00% | 0.00000 | 0.420160 |
| Two Patches (Transfer learning) (pool5 - layer8) - 32 batches | 92.00% | 0.04470 | 8.00% | 84.00% | 0.08940 | 86.67% | 0.07450 | 100.00% | 0.00000 | 0.406120 |
| Two Patches (Transfer learning) (pool5 - layer9) - 32 batches | 94.00% | 0.05480 | 6.00% | 88.00% | 0.10950 | 90.00% | 0.09130 | 100.00% | 0.00000 | 0.420160 |

Additional testing was carried out in order to confirm that the CNN architecture is not biased due to color space, for the ImageNet dataset is RGB while our dataset is intrinsically grayscale. Hence, we retrained the network with 200 grayscale images and reapplied our investigation. The result was conclusive that the network is independent of the color space as the results obtained exactly matched that of directly using the ImageNet pretrained CNN.

Significant degradation of performance resulted upon downsampling, as shown in Table 1. After training with downsampled images, accuracy was at most 78% (71% specificity, 78% precision, and 100% recall). Every metric, except for the recall, was lower comapred to that of the CNN trained at full resolution. In conclusion, resampling of input should be avoided, and extraction of patches as per the proposed methodology is recommended.

Finally, the confusion matrix of the chosen CNN for the proposed early DR detection CAD system is shown in Fig 9. A comparison of our proposed technique against other machine learning techniques shows its superiority as can be observed in Table 2.

**Table 2.** Comparison of our proposed convolutional neural network (CNN) with four other machine learning classifiers.

| Classifier | Accuracy | Recall | Specificity |
| --- | --- | --- | --- |
| *CNN (proposed)* | 94% | 100% | 88% |
| *K-Star(\*)* | 89% | 89% | 89% |
| *K-Nearest Neighbor (kNN)* | 84% | 84% | 83% |
| *Random Forest* | 85% | 82% | 82% |
| *Random Tree* | 81% | 81% | 81% |

**Fig 9.**
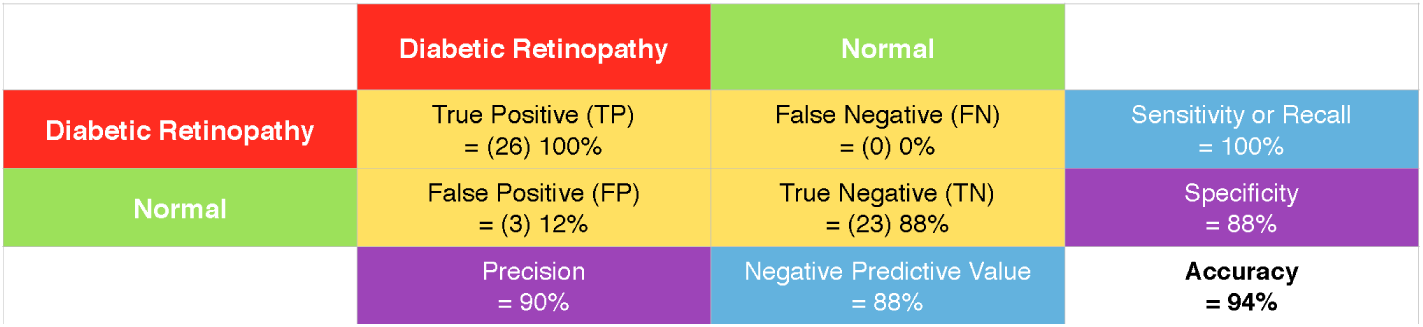
Confusion matrix for proposed computer aided design system for early DR detection with CNNs.

## Conclusion

Early intervention is essential to delay or prevent complications of diabetic retinopathy, including blindness. As such, in this paper, a novel CAD system for early detection of DR-related changes in OCT images using CNNs was presented. The system was developed for use patients with almost clinically normal retina appearances.

Upon investigation of the various proposed parameters of the CAD system, the optimal conditions for its deployment were derived. Foremost, transfer learning may be used to achieve high accuracy despite the scarcity of the data. Second, best results are seen when combining the output features of two independently trained CNN, which operate on either side of the fovea. Features extracted at the *pool5* layers of these CNN provided for the highest accuracy with the least computational complexity. Finally, in order to reach highest accuracy, which our preliminary results found to be 94 %, the patches extracted for training and testing should be aligned or centered along the *y*-axis using the patch extraction algorithm presented and the segmented OCT layer number 6, or the outer nuclear layer (ONL). This paper recommends that further research is directed towards this relatively uncharted topic, especially with OCT images, for the results were observed to be high even given the scarcity of the data and the relative complexity of the problem.

